# Transcriptional and Metabolic Mechanism of Carbon Dots Enhancing Rice Growth and Resistance by Promoting Root

**DOI:** 10.1101/2024.08.01.606230

**Authors:** Yadong Li, Ronghua Xu, Jingyi Qi, Shang Lei, Qianying Han, Congli Ma, Yunlong Ru, Hongjie Wang

## Abstract

Increasing climate change and pollutant discharge induce constant challenges to crops, while crops are vulnerable to environmental and pollutant stresses. In this study, a carbon dots (CDs) was developed that significantly increased rice seedling growth, and successfully reduced the inhibition of heavy metal cadmium (Cd), salt (NaCl), and herbicide 2,4-D stresses on rice seedling growth by pre-spraying. The root of rice seedlings responded specifically to CDs exposure, with significant improvements in root biomass, architecture, cell wall thickness, mechanical strength, and metabolic vitality. Metabolomics and transcriptomics were combined to reveal the regulatory mechanism of CDs in rice seedlings. Transcriptome analysis indicated that CDs upregulated genes related to cytokinin, jasmonic acid, salicylic acid, MAPK signaling pathway, calcium homeostasis, and peroxidase, and downregulated those related to auxin, abscisic acid, and ethylene. Metabolomic analysis suggested CDs improved the metabolites related to antioxidant (betalain, ascorbate, aldarate, and glutathione), formation of cell wall, plasma membrane, xylem, and root cortex (phenylpropanoids biosynthesis, stilbenoid, diarylheptanoid and gingerol biosynthesis, and sphingolipid), and energy metabolism (nicotinate, nicotinamide, glyoxylate, dicarboxylate, and nitrate cycle) in rice seedlings. Therefore, pre-spraying CDs reprogrammed stress signaling pathways and enhanced adaptive responses in rice seedlings, ultimately increasing growth potential and stress resistance. This study presents a promising nano-bio-stimulant of CDs for crop resilience in the context of increasing climate change and contributes to sustainable agriculture.

## INTRODUCTION

“Zero hunger” is the goal of sustainable agriculture. Yet, over 258 million people worldwide still face severe food insecurity in 2022 (FAO et al., 2023). Food production urgently needs to increase, however, agriculture faces growing challenges from climate change, soil degradation, and environmental contamination. Abiotic factors, such as drought, heat, cold, nutrient deficiencies, and salinity, are fundamental environments that adversely affect plant distribution and productivity (Zhu, 2016; Zhang et al., 2022). Rapid industrial development and the misuse of agrochemicals lead to significant pollution in agricultural soils, including heavy metals and pesticide residues (Riedo et al., 2021; Fei et al., 2022). With staggering input of resources, the yield curves of many crops are flattening (Kah et al., 2019; Lowry et al., 2019). These situations call for a new agri-tech revolution to sustainably enhance crop stress resistance in response to the increasing global demand for food.

Plants have evolved sophisticated and interconnected immune systems to defend against various adverse environments. Adverse environments disrupt the biomolecules and metabolic processes in plants, leading to fluctuations in second messenger levels such as Ca^2+^, reactive oxygen species (ROS), nitric oxide, and phospholipids (Zhu, 2016). These changes will trigger stress-specific signal transductions and adaptive/resistant responses in plants (Zhang et al., 2022). Ca^2+^ signaling has been recognized as the front line of translators for different environmental stresses due to its rapid response within seconds (Ku et al., 2018). Additionally, phytohormones in plants account for long-distance signaling over different organs. Especially, abscisic acid (ABA), jasmonic acid (JA), and salicylic acid (SA) are major phytohormones that channel resources into mitigating abiotic stresses (Ku et al., 2018). Therefore, it can be proposed that pre-stimulating the immune system of plants before environmental stress could significantly reduce the adverse influence.

Advances in engineered nanomaterials are attracting increasing attention in agriculture (Lowry et al., 2019; Usman et al., 2020). The small size (<100 nm) and high specific surface area enable nanomaterials to easily penetrate the biological barriers and actively interact with cellular components. Recently, carbon dots (CDs, size about 3~5 nm), with physical and chemical properties, good biocompatibility, and low toxicity, have shown significant potential in biomedical, environmental, and agricultural applications (Peng et al., 2017; Ghosal and Ghosh, 2019; Long et al., 2021). CDs are commonly reported to promote plant photosynthesis due to the photoluminescence and improvement of the photosynthetic activity (Li et al., 2018; Li et al., 2021; Li et al., 2021). Moreover, CDs also contribute to plant resistance against abiotic/biotic stresses, including drought (Su et al., 2018; Yang et al., 2022), salinity (Li et al., 2022), heavy metals (Xiao et al., 2019), and diseases (Li et al., 2018), which usually are simply attributed to the radical scavenging of CDs and enhancements of the antioxidant defense system of plants. However, in our previous study, pre-treatment of CDs significantly enhanced the resistance of rice seedlings when exposed to abiotic stresses (Li et al., 2020). We proposed that pre-treatment of CDs might act as a nano-bio-stimulant to trigger the immune system or improve the growth potential of plants, thereby enhancing stress resistance.

Herein, we developed a CDs and found that pre-spraying CDs at suitable doses significantly enhanced the resistance of rice seedlings to salt, heavy metal cadmium (Cd), and herbicide 2,4-Dichlorophenoxyacetic acid (2,4-D) stresses. Especially, the roots of rice seedlings showed meaningful responses to CDs exposure. Subsequently, integrated analysis of transcriptome and metabolome was employed to elucidate the regulatory mechanism of CDs in rice seedlings. The study provides a new and systematic understanding of CDs improving plant stress resistance contributes to the nano-enabled agri-tech revolution.

## RESULTS

### Characterization of CDs

The prepared CDs exhibited blue fluorescence with maximum light absorption, fluorescence excitation, and emission peaks centered at 320 nm, 360 nm, and 430 nm, respectively (Figure 1a and Figure S2A). The fluorescence emissions of the CDs were obviously excitation-dependent (Figure S2B). TEM images (Figure 1b) show that the spherical CDs particles are well monodispersed with an average size of 2.61 nm and a lattice spacing of 0.20 nm, corresponding to (101) facet of graphite. Figure 1c shows the FTIR spectrum of CDs, with stretching vibration of the NH/−OH bonds in −NH_2_, −COOH, or −OH groups around 3400 cm^−1^, the C=O bond in ketone group around 1713 cm^−1^, the C=O/N-H bond in amide/amine groups around 1636 cm^−1^, and the C-O/C-N bonds around 1405, 1339, and 1219 cm^−1^. XPS survey spectrum indicates that the CDs are composed of C (284.8 eV), O (400.2 eV), and N (532.3 eV) elements with atomic ratios of 55.73%, 41.58%, and 2.7%, respectively (Figure 1d). C1s spectrum was deconvoluted into C–C, C–O–C, and O–C=O bonds at 284.8 eV, 286.6 eV, and 288.9 eV, respectively (Figure 1e) (Lan et al., 2017). The high-resolution spectrum of O1s reveals the C-O bond at 531.8 eV and the C=O bond at 532.8 eV (Figure 1f), while that of N1s shows graphitic N at 400.4 eV (Figure 1g). Therefore, it can be inferred that the CDs is covered with a highly hydrophilic surface containing −NH_2_, −COOH, −OH, and −CONH_2_ groups, which significantly contribute to its uptake and interaction with biological systems. The n-π* transitions of the p-π orbit between the organic moieties on the surface and the conjugate structure of the CDs are responsible for photon capture, electron transitions, and fluorescence emission (Sun et al., 2015; Zhang et al., 2016).

**Figure 1.**
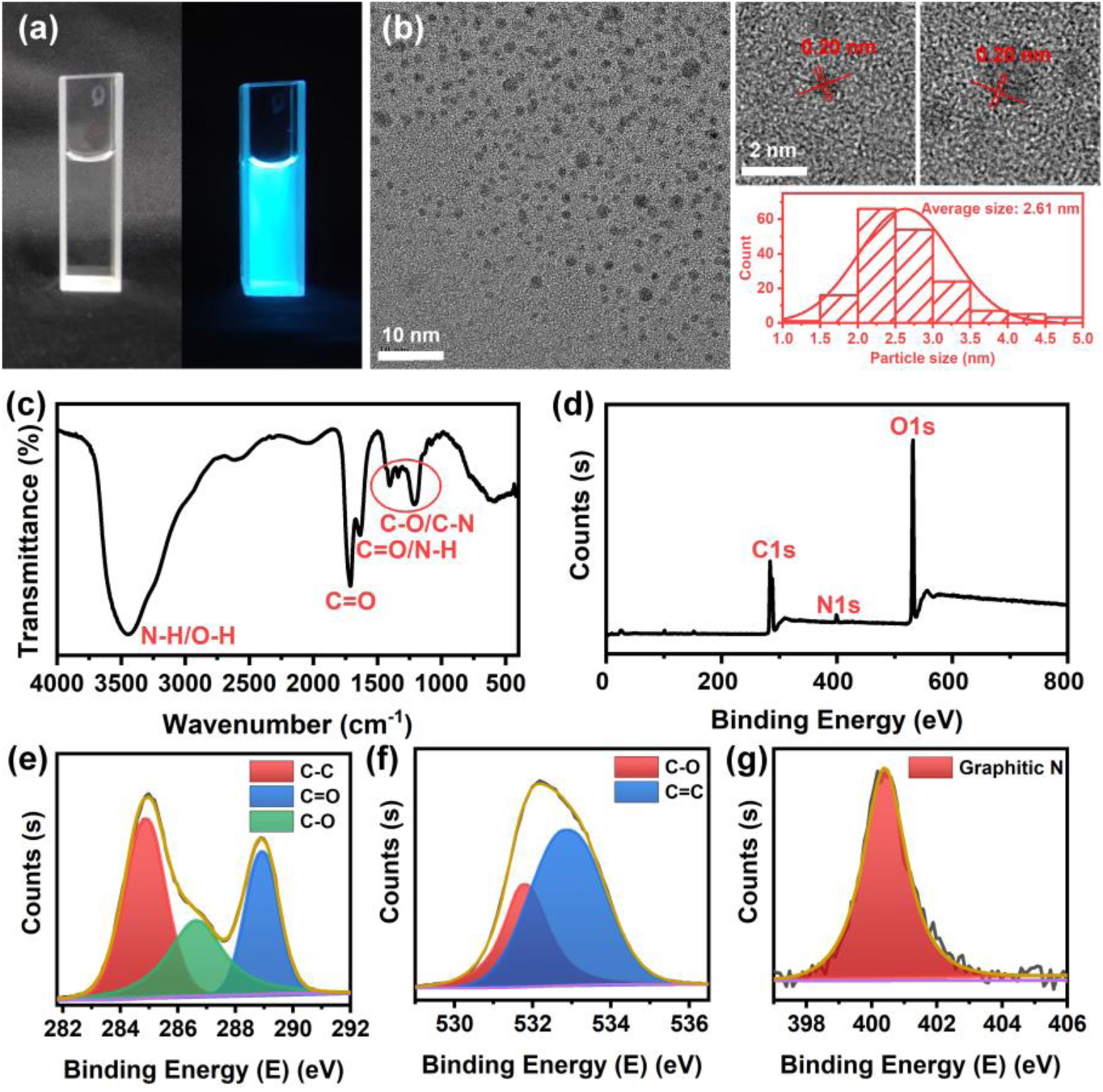
Morphology, chemical structure, and component of the prepared CDs: (a) photograph of CDs under daylight and 365 nm light; (b) TEM images, lattice fringe, and size distribution histograms of CDs; (c) FTIR spectrum of CDs; (d) full XPS patterns and (e-g) high-resolution XPS spectra of C1s, O1s, N1s peaks of CDs.

### CDs enhanced rice growth and resistance to abiotic stresses

Without stresses, spraying CDs significantly increased the fresh biomasses of rice seedlings in a concentration-dependent manner rather than the elongation of rice roots and shoots. The maximum improvements of 53.04%, 33.56%, and 39.06% in root, shoot, and total fresh weights were induced by CDs at 100 μg/mL (Figure S3A and B). CDs at 100 μg/mL also significantly increased the chlorophyll index of rice seedlings by 8.67% (Figure S3C). It seems that foliage spraying of CDs is preferred to promote root development (Figure S3D).

Under Cd and NaCl stresses, the root length, root surface area, root volume, leaf area, leaf length, and fresh weight of CDs-pre-sprayed rice seedlings were significantly increased by 21.05%, 42.38%, 101.05%, 34.51%, 12.77%, 48.20% (Cd stress), and 45.50%, 97.96%, 206.03%, 143.63%, 31.85%, 116.80% (NaCl stress) compared to those without CDs exposure, respectively. For 2,4-D stress, pre-spraying of CDs significantly increased the root surface area and volume of rice seedlings by 55.42% and 92.87%, respectively (Figure 2 and Figure S4). Cd, NaCl, and 2,4-D stresses induced obvious leaf scorch, browning, and necrotic lesions on rice leaves, which were not observed on those pre-sprayed with CDs. These results suggested that pre-spraying CDs improved the growth potential and resistance prior to the environmental stresses, and the response of root may play key roles.

**Figure 2.**
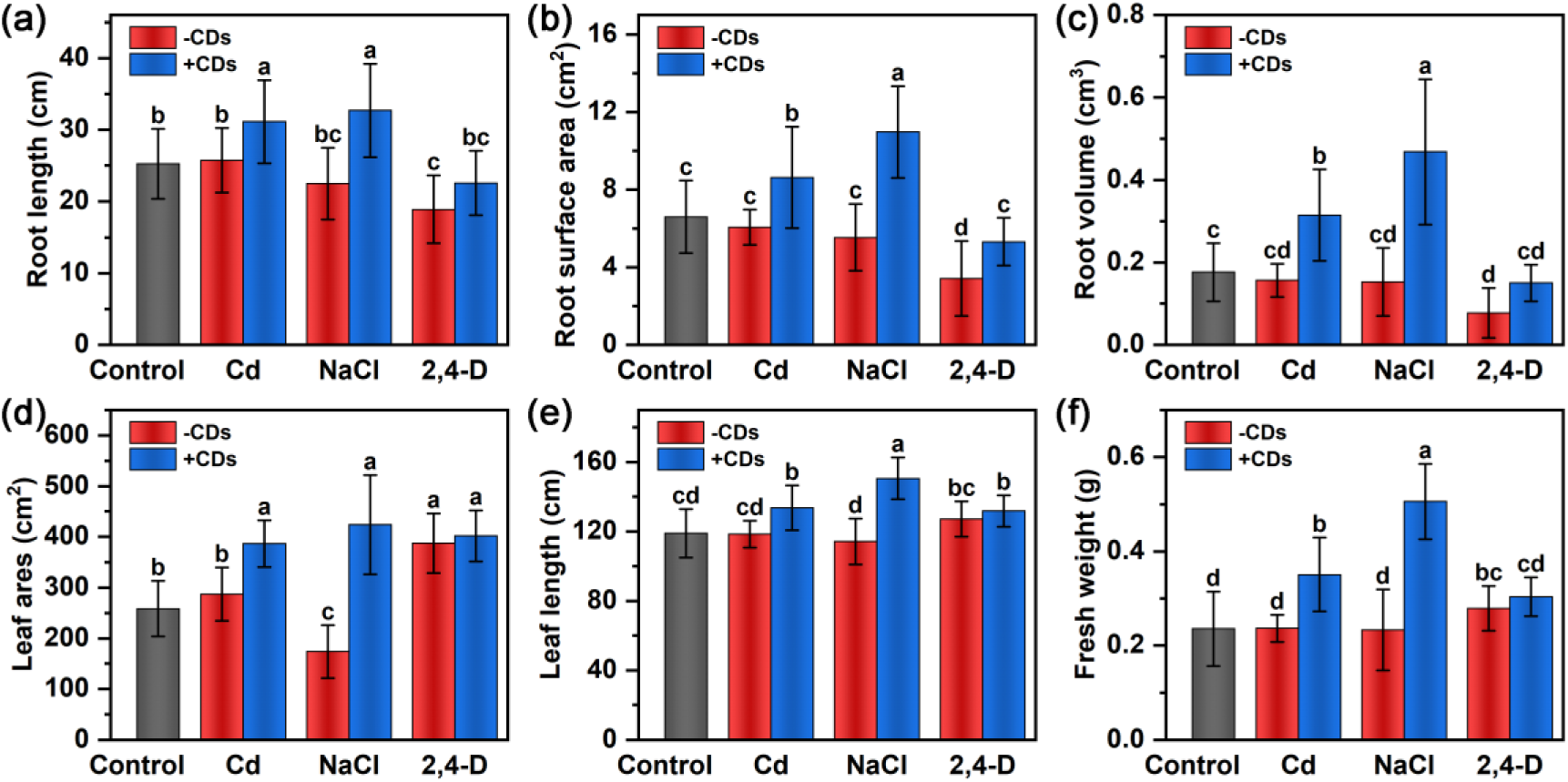
Pre-spraying of CDs (100 μg/mL) increased rice seedling growth under heavy metal Cd, salt, and herbicide 2,4-D stresses: (a) root length, (b) root surface area, (c) root volume, (d) leaf area, (e) leaf length, and (f) fresh weight. Error bars correspond to standard deviation (n ≥ 10). Marked with asterisk (*) indicates a significant difference (*p* < 0.05).

### CDs improved root development and vitality of rice seedlings

Then, the root development, anatomy, microstructure, and vitality of CDs-pre-sprayed rice seedlings were further. TEM images (Figure S5) show that CDs are widely distributed in both the leaves and roots of rice seedlings, including cell walls, chloroplasts, and cytoplasm. As shown in Figure 3a and b, rice roots pre-sprayed with CDs are significantly stronger than those in the control group, with total length, surface area, volume, and diameter being increased by 9.44%, 29.66%, 18.63%, and 16.98%, respectively. The paraffin sections of rice roots stained with Safranine O/Fast Green (Figure 3c) indicate that CDs increased the lignocellulosic constituents in the epidermis, cortex, and vascular cell walls due to more tissues in green. SEM images (Figure 3d) show the strengthened root surface of rice seedlings by CDs with much fewer breakages. Besides, CDs-pre-sprayed rice roots exhibit more irregular or elongated cells, thicker cell walls, and more cell contents compared to those in the control group (Figure 3e). The CDs-thickened cell wall is also observed in rice leaf cells (Figure S5). Furthermore, the root vitality, cytoplasm content, and concentrations of Ca, Mg, and Mn ions were significantly increased by 115.09%, 39.52%, 9.37%, 13.07%, and 19.29% in CDs-pre-sprayed rice seedlings, respectively (Figure 3f-h). These results suggest that the sprayed CDs particles permeated into rice mesophyll cells and then were transferred to roots. In rice seedlings, CDs enhanced the secondary cell walls, mechanical defense, supporting anchorage, metabolic vitality, and cell division of rice roots, which finally increased rice growth and resistance to stresses.

**Figure 3.**
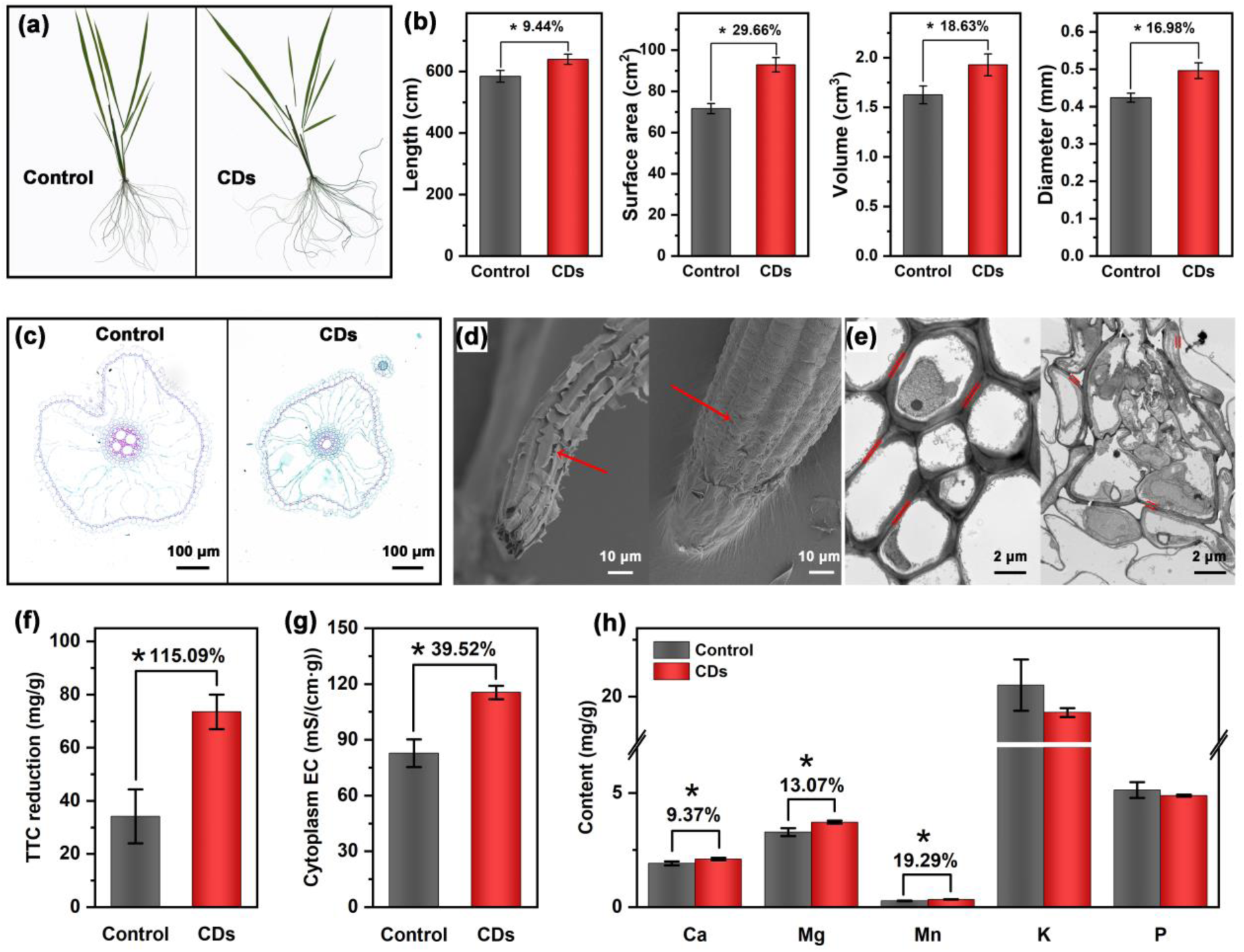
Pre-spraying of CDs (100 μg/mL) improved rice root development, mechanical strength, and cell metabolism: (a) scan images of rice seedlings; (b) length, surface area, volume, and diameter of rice roots; (c) paraffin section images of rice roots stained by Safranine O/Fast Green; (d) SEM and (e) TEM images of rice roots in control and CDs treatments, red arrows and lines indicate the breakages on root surface and cell wall thickness, respectively; (f) root vitality, (g) electrical conductivity (EC) of cytoplasm content; (h) Ca, Mg, Mn, K and P contents. Error bars correspond to standard deviation (n ≥ 3). Marked with asterisk (*) indicates a significant difference (*p* < 0.05).

### Transcriptional changes in CDs-pre-sprayed rice seedlings

The transcriptome of CDs-pre-sprayed rice seedlings was further measured to uncover the molecular events responsible for the enhanced stress resistance of rice seedlings. A total of 23833 genes were detected from all rice seedlings, of which 1062 and 720 genes were specifically expressed in the control and CDs treatments, respectively (Figure S6A). Principal component analysis (PCA) revealed the CDs-changed transcriptome profile in rice seedlings, with 1026 upregulated and 858 down-regulated DEGs (differentially expressed genes) (FC>2 or FC<0.5, *p*≤0.05) (Figure S6B and C). These DEGs enriched in 7 KEGG pathways, including diterpenoid biosynthesis, phenylpropanoid biosynthesis, plant hormone signal transduction, MAPK signaling pathway-plant, plant-pathogen interaction, cysteine and methionine metabolism, and zeatin biosynthesis (Figure 4a). Notably, all these pathways are associated with stress and immune systems in plants. As shown in Figure 4b, apart from various GO terms related to intracellular redox balance and resistance to oxidative stress, a maximum of 130 genes enriched in GO term of transmembrane transport. Thus, it can be inferred that CDs pre-triggered the immune system of rice seedlings prior to the environmental stresses.

**Figure 4.**
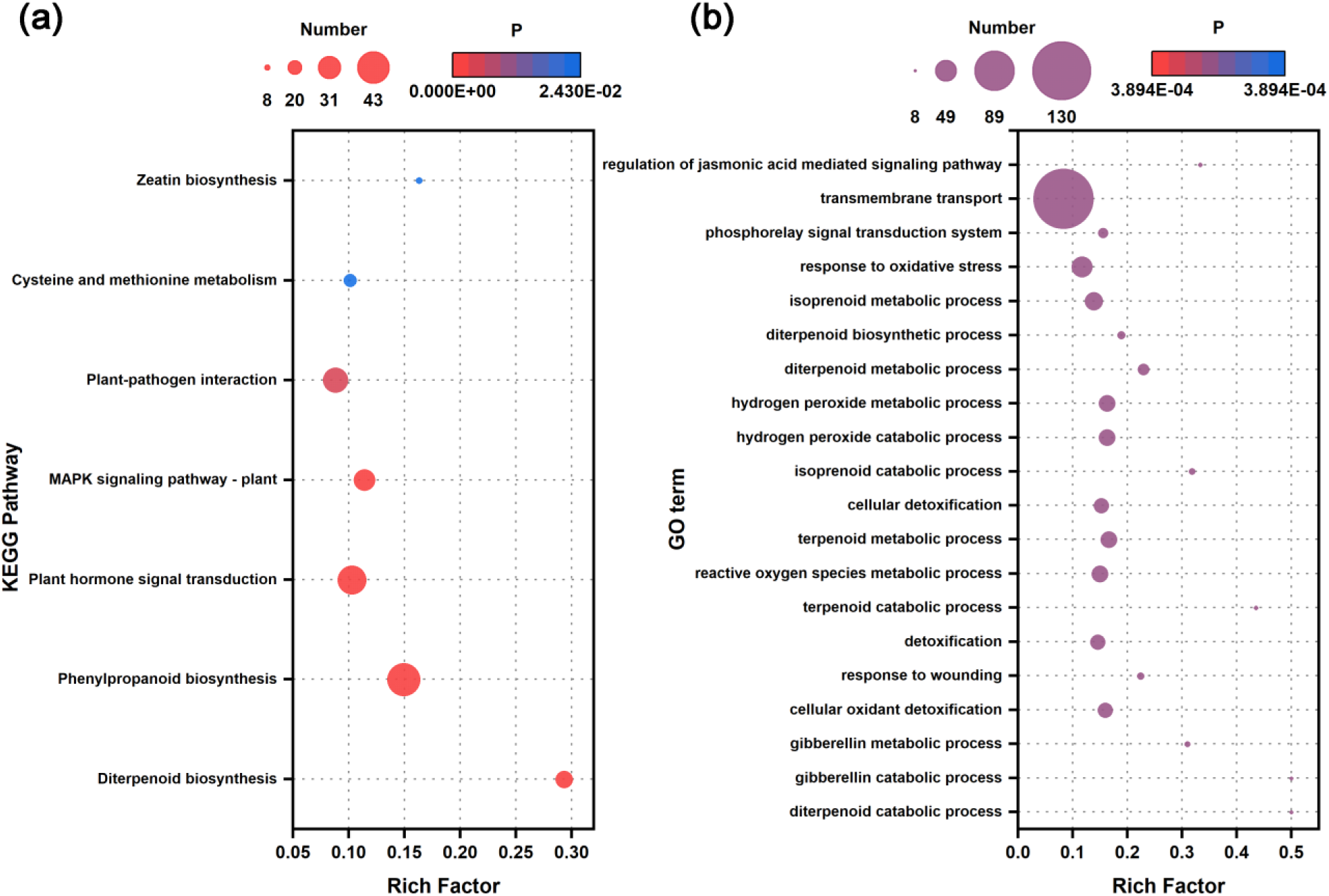
Bubble diagram of (a) GO and (b) KEGG enrichment analysis of DEGs in rice seedlings induced by CDs, only showing the top 20 enriched GO term/KEGG pathways with *p*-value≤0.05. Note: the vertical axis represents the GO term/KEGG pathway, the horizontal axis represents the rich factor, and the size and color of the point represent the number of genes in the corresponding GO term/KEGG pathway and different *p* ranges, respectively.

In plant hormone signal transduction pathway, CDs induced phytohormone-dependent regulation in rice seedlings (Figure 5a). The genes related to auxin, abscisic acid, ethylene, and their signaling cascades were down-regulated by CDs, while those of cytokinin, jasmonic acid, and salicylic acid were up-regulated. In the MAPK signaling pathway, pre-spraying of CDs up-regulated the genes associated with flg22 and H_2_O_2_, which mediated plant resistance to pathogen attack and infection. Especially, the up-regulated genes of MEKK1 and MPK3/6 are widely involved in enhancing the tolerance of plants to both biotic and abiotic stresses (Figure 5b). As responses, pre-spraying of CDs significantly up-regulated genes encoding calcium-dependent protein kinases (CDPKs) and calmodulin-like proteins (CMLs) in plant-pathogen interaction pathway (Figure 6a), as well as peroxidase (POD, EC: 1.11.1.7) in phenylpropanoid biosynthesis pathway (Figure 6b). In comparison, the CDs-induced DEGs of rice showed obvious down-regulation trends in the diterpenoid biosynthesis pathway and irregular regulations in cysteine and methionine metabolism and zeatin biosynthesis pathways (Figure S7). In the antioxidant defense system of rice seedlings, the enzymatic activity of POD was significantly increased by 160.74% in CDs pre-sprayed treatment, the superoxide dismutase (SOD) and catalase (CAT) activities were significantly decreased by 20.52% and 37.60%, and the malondialdehyde (MDA) content was not significantly affected (Figure S8). Therefore, pre-spraying of CDs on rice seedlings not only improved jasmonic acid, salicylic acid, and MAPK pathways to signal environmental stresses but also enhanced Ca^2+^ homeostasis and peroxidase-mediated antioxidant activity to tolerate environmental stresses.

**Figure 5.**
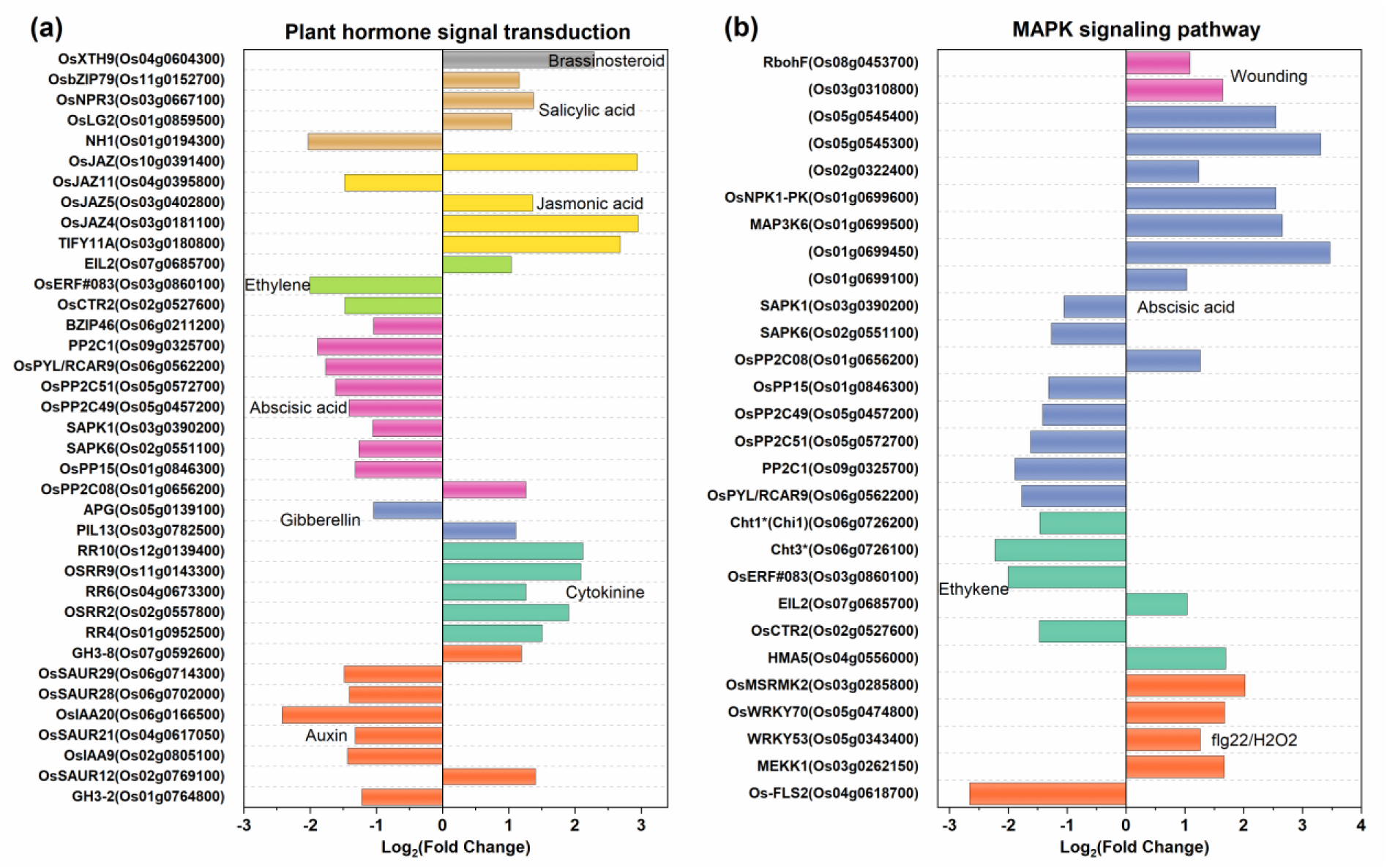
Gene expression levels of DEGs enriched in (a) plant hormone signal transduction and (b) MAPK signaling pathways.

**Figure 6.**
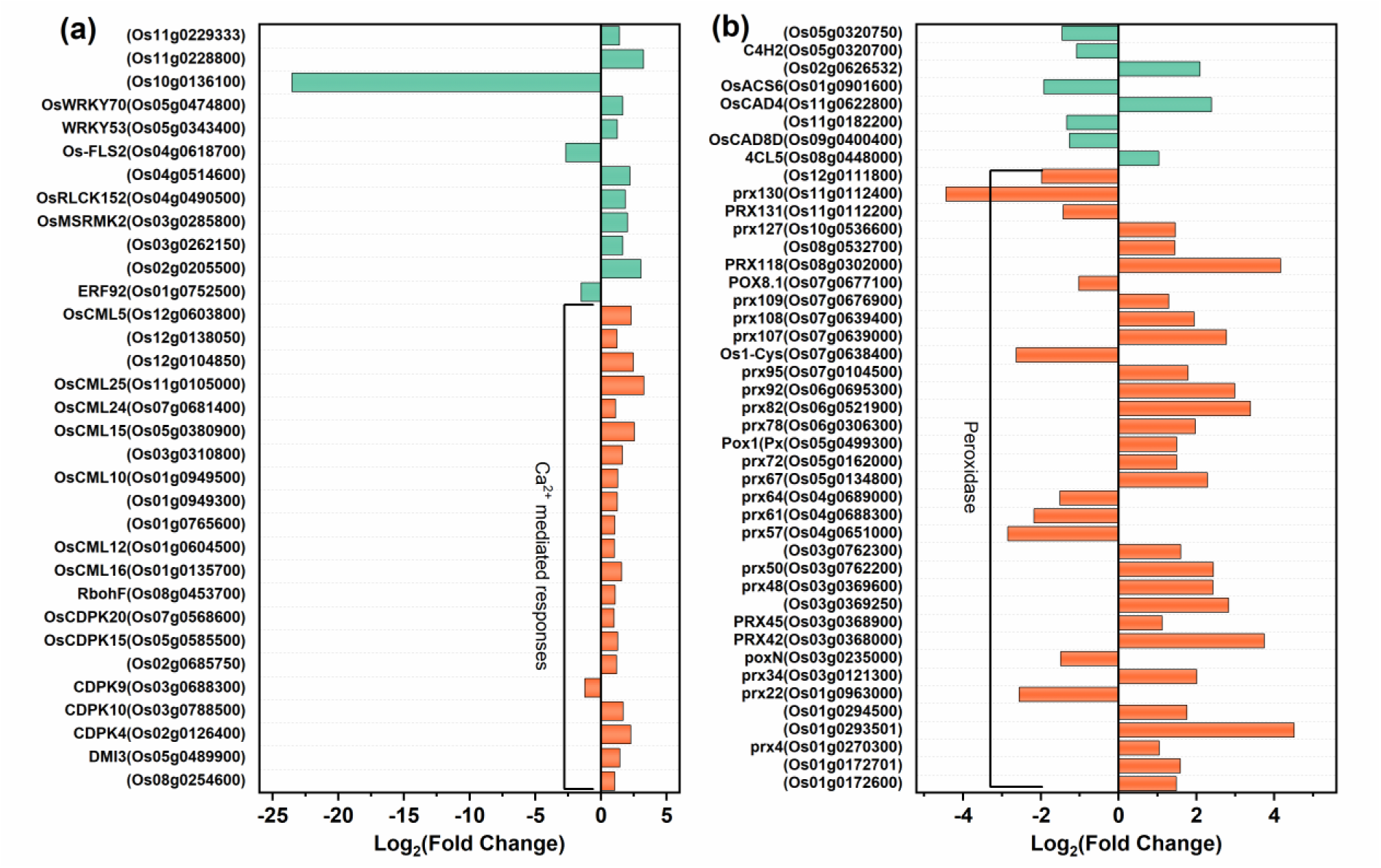
Gene expression levels of DEGs enriched in adaptability-related KEGG pathways in CDs-treated rice seedlings: (a) plant-pathogen interaction, (b) phenylpropanoid biosynthesis.

### Metabolomics changes in CDs-pre-sprayed rice seedlings

Transcriptome analysis revealed the changes in stress defense-related genes induced by pre-spraying CDs in rice seedlings. Then, the untargeted metabolomics of the CDs-pre-sprayed rice seedlings was further profiled to determine the relative abundance of metabolites in rice seedlings. A total of 2274 metabolites were identified in rice seedlings, with 84 and 69 metabolites being significantly up- and down-regulated by pre-spraying CDs (FC>1, *p*≤0.05) (Figure S9A and B). PCA plot indicates that pre-spraying induced significant alternation in the transcriptome profile of rice seedlings (Figure S9C). In KEGG enrichment analysis of the DEMs, most of the significantly enriched pathways show up-regulations, including lysine degradation, nicotinate and nicotinamide metabolism, citrate cycle (TCA cycle), ascorbate and aldarate metabolism, phenylpropanoid biosynthesis, nucleotide metabolism, betalain biosynthesis, glyoxylate and dicarboxylate metabolism, stilbenoid, diarylheptanoid and gingerol biosynthesis, sphingolipid metabolism, and glutathione metabolism. While the pathways of flavonoid biosynthesis, C5-branched dibasic acid metabolism, and arginine biosynthesis were down-regulated by CDs (Figure 7).

**Figure 7.**
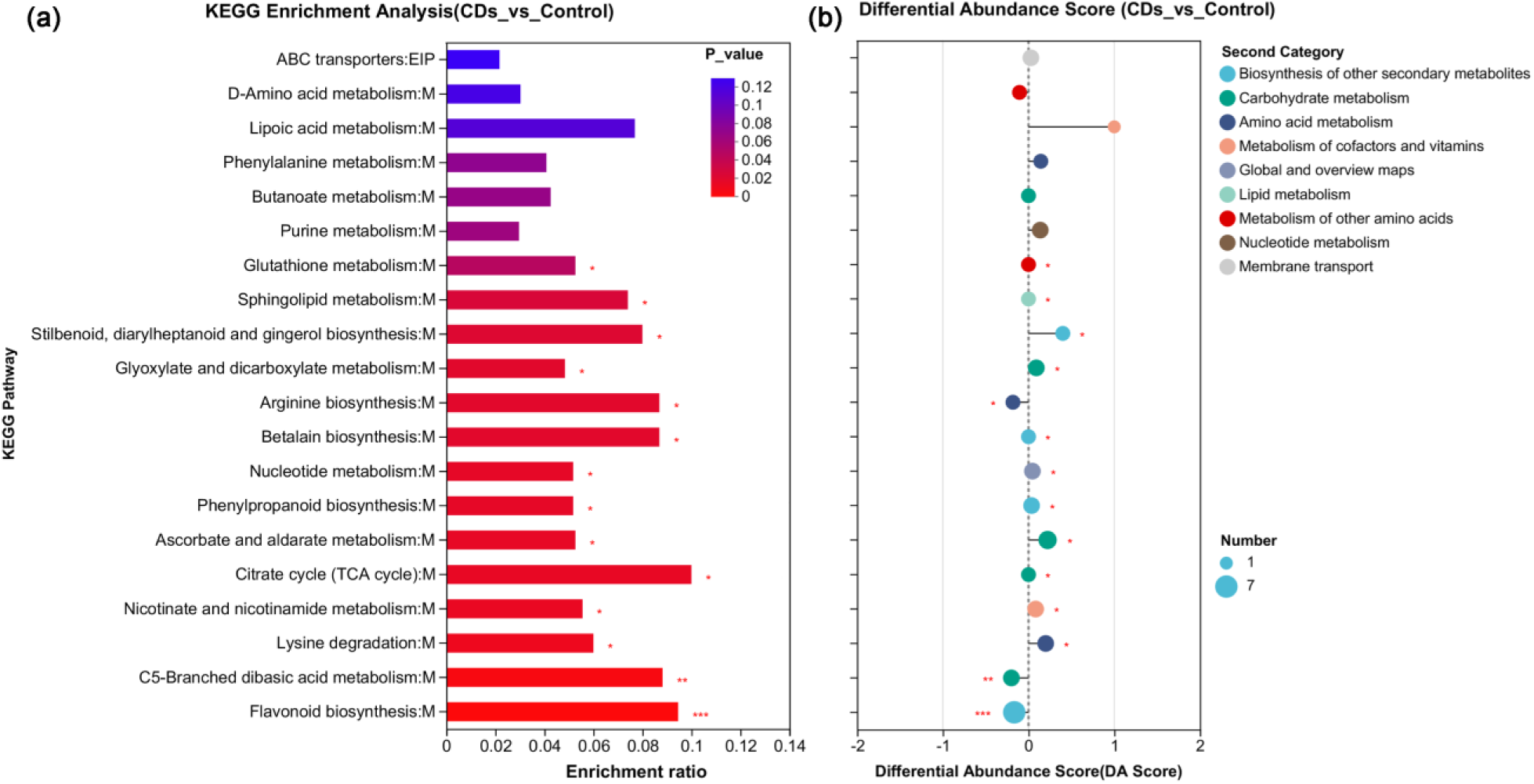
(a) KEGG enrichment analysis and corresponding (b) differential abundance score of DEMs in the metabolome of rice seedlings induced by CDs. Only showing the top 20 enriched KEGG pathways with *p*-value≤0.5, marked with one, two, and three asterisks (*) indicates a significant difference with *p* < 0.05, *p* < 0.01, and *p* < 0.001, and the dots on the right of the medial axis in (b) mean upregulations and on the left mean downregulations.

Lysine content in plants show widely negative correlations with endosperm development, seed germination, starch synthesis, and grain yield in various plants (Yang et al., 2020). High free lysine content also can attenuate the tricarboxylic acid (TCA) cycle by affecting the related metabolites and gene transcript (Angelovici et al., 2011). Therefore, the up-regulated lysine degradation by pre-spraying CDs, alone with nucleotide metabolism, contributed to the growth and development of rice seedlings. Besides, the up-regulations of betalain, ascorbate, aldarate, and glutathione, the key metabolites with strong antiradical and antioxidant activities in plants, could significantly protect plant cells and tissues against excess ROS accumulations. In addition, the pathways of phenylpropanoids biosynthesis, stilbenoid, diarylheptanoid and gingerol biosynthesis, and sphingolipid metabolism participate in the formation of cell walls, plasma membrane, xylem, and root cortex, which contribute to anchoring and supporting plants as well as the absorption and transport of nutrients (Quinville et al., 2021; Rahim et al., 2023). Furthermore, the nicotinate and nicotinamide metabolism, nitrate cycle (TCA cycle), and glyoxylate and dicarboxylate metabolism are the most important energy metabolism pathways. The combination of the effects of pre-spraying CDs on the metabolomics of rice seedlings would significantly improve the growth and development, biochemical and histochemical resistance to biotic and abiotic stresses.

## DISCUSSION

With excellent photoluminescence, CDs attracted most attentions on plant photosynthesis (Li et al., 2018; Li et al., 2021; Li et al., 2021). Some effort of CDs has been made to increase plant resistance against abiotic stresses, which was often attributed to the radical scavenging activity of CDs and their improvement to plant antioxidant defense system (Su et al., 2018; Xiao et al., 2019; Yang et al., 2022). In this study, we found pre-spraying CDs could significantly protect rice seedlings from environmental stresses by pre-triggering the immune system and adaptive responses. According to the combined analysis of transcriptomics and metabolomics, the enhancement of rice resistance by pre-spraying CDs can be attributed to the improved stress signaling, nutrition assimilation and metabolism, biochemical and mechanical defenses of rice seedlings, which endowed rice with higher growth potential and resistance when exposed to stresses.

At the upstream of gene transcription level, prs-spraying CDs significantly up-regulated genes related to jasmonic acid, salicylic acid, and MAPK signaling pathway, which improved the capacity of rice seedling to sense and signal environmental stress (Pieterse et al., 2012). The other up-regualtions to cytokinin-related genes and down-regulations to auxin-, abscisic acid-, and ethylene-related genes in rice seedlings would lead to a complicated network to determine the root development and secondary cell wall: auxin and cytokinin antagonistically define the root apical meristem size; Jasmonate and abscisic acid antagonistically cooperated with auxin to determine the root apical meristem activity; Ethylene and auxin mutually interconnect to control the rapid elongation of the differentiated cells at the periphery of the root apical meristem (Vanstraelen and Benková, 2012); The crosstalk between brassinosteroids, cytokinin, auxin, jasmonate, and abscisic acid determine the xylogenesis and secondary cell wall formation by regulating the transcriptional cascade of related genes; And the brassinosteroids, auxin, cytokinin, ethylene, and gibberellins are involved in regulating genes of cellulose, xylan biosynthesis, and lignification in secondary cell wall structure (Didi et al., 2015). Furthermore, metabolomics of rice seedlings showed upregulations in phenylpropanoids biosynthesis, stilbenoid, diarylheptanoid and gingerol biosynthesis, and sphingolipid metabolism pathways by pre-spraying CDs for the formation of cell walls, plasma membrane, xylem, and root cortex. Therefore, these alternations induced by pre-spraying CDs finally enhanced the stress sensing and signaling, root development, and cell wall reinforcement of rice seedlings.

Moreover, pre-spraying CDs up-regulated genes encoding CDPKs, CMLs (control the Ca^2+^ homeostasis in plant cells), and peroxidase as well as metabolites in pathways of betalain biosynthesis, ascorbate and aldarate metabolism, and glutathione metabolism in rice seedlings. At the physiological and biochemical levels, pre-spraying CDs increased Ca, Mg, and Mn element contents and POD enzymatic activity in rice seedlings. Thereby, these changes induced by pre-spraying CDs contributed to the Ca^2+^ homeostasis and signaling abiotic stress (Schulz et al., 2013; Zeng et al., 2015), antiradical and antioxidant activities, and mineral nutrient assimilations, which ensured the ion, osmosis, and REDOX homeostasis of the intracellular environment in rice seedlings when exposed to stresses. The upregulated CMLs genes also participate in cell cycle control, cell division, chromosome partitioning, and cytoskeleton (Table S2). Furthermore, pre-spraying CDs also led to significant enhancement to the nicotinate and nicotinamide metabolism, citrate cycle (TCA cycle), and glyoxylate and dicarboxylate metabolism pathways. TCA cycle is a universal pathway in mitochondria to oxidate respiratory substrates in plants, producing sufficient energy for life activities of plants and intermediates of organic acids for biosynthesis of amino acids, fatty acids and secondary metabolites (Tavsan and Ayar Kayali, 2015), Glyoxylate and dicarboxylate metabolism in glyoxylate cycle act as a branch of the TCA cycle to produce energy when the intermediates are insufficient. Nicotinate and nicotinamide in plants can be converted into nicotinic acid adenine dinucleotide (NAD) and nicotinamide adenine dinucleotide (NADP), which act as coenzymes to participate in the energy metabolism and biochemical reactions. Therefore, pre-spraying CDs promoted the energy metabolism of rice seedlings, thereby imrpoving the metabolic and growth activity.

In conclusion, this study developed a strategy of pre-spraying CDs as a novel nano-biostimulant to enhance crop growth and resistance against abiotic stresses. The underlying regulatory mechanism of CDs in rice seedlings was showed in Figure 8. Firstly, after spraying, CDs permeated into mesophyll cells and were transferred to roots *via* the vascular system. In rice seedlings, CDs triggered stress/immune system and induced adaptive responses by reprogramming phytohormone and MAPK signaling pathway, Ca^2+^ homeostasis, antioxidant-related enzymes and metabolites, nutrition assimilation, and energy and nucleotide metabolism. These regulations of CDs subsequently induced enhancements in root development and vitality for nutrient acquisition and metabolism, mechanical strength of cell walls and root to defense environments, REDOX homeostasis, and metabolic activity in rice seedling when exposed to stress. This study presents an ideal strategy for agriculture in response to escalating climate change and contributes to the nano-enabled agri-tech revolution for sustainable agriculture.

**Figure 8.**
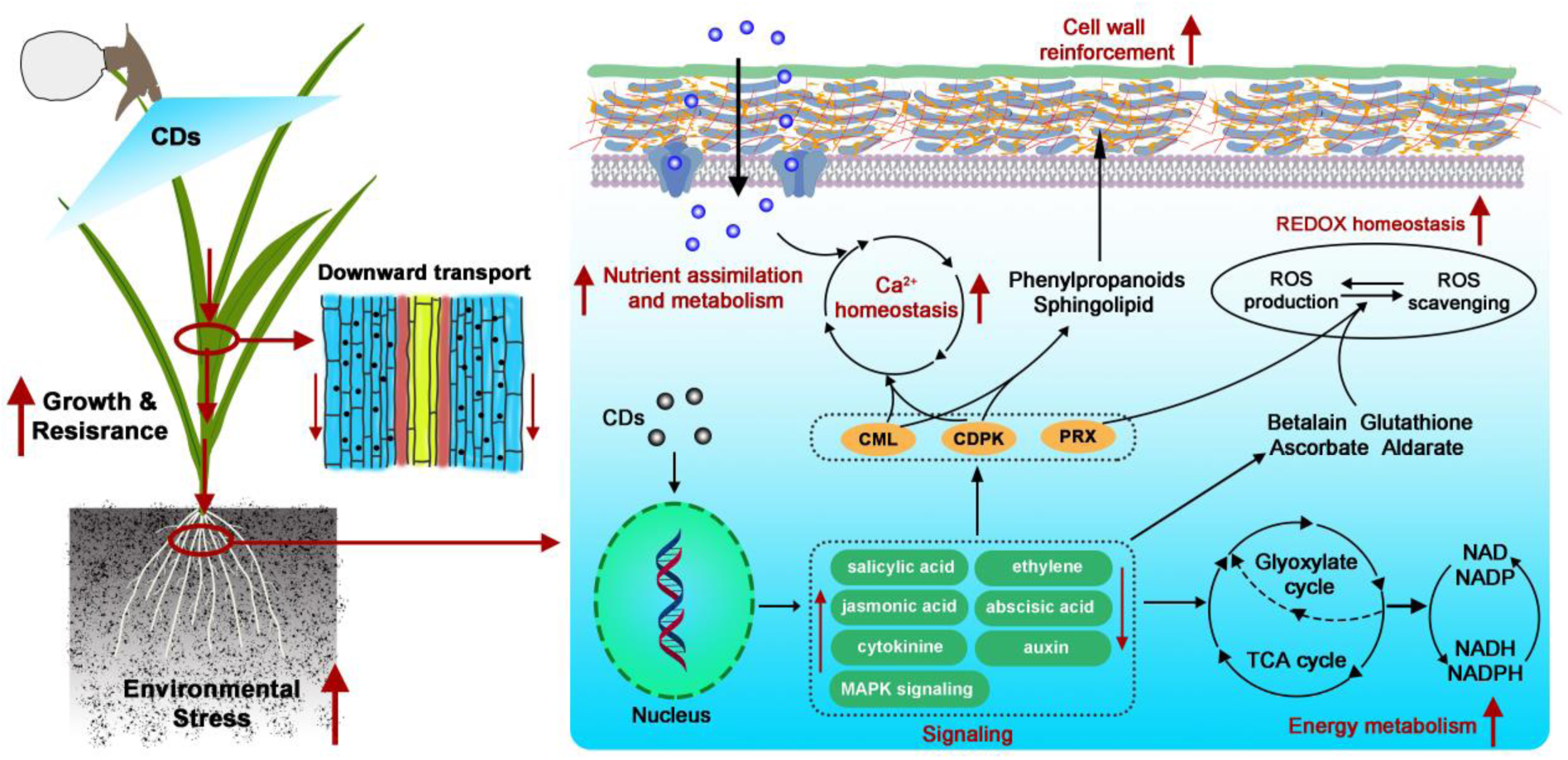
Schematic representation of the mechanism CDs to pre-stimulate rice growth and stress resistance. CDs in rice reprogrammed the phytohormones and MAPK signaling pathways, which led to significant enhancements to cell wall reinforcement, nutrient assimilation, Ca^2+^ homeostasis, REDOX homeostasis, and energy metabolism of rice seedlings, thereby improving rice growth and stress resistance. CML: calmodulin-like proteins; CDPK: calcium-dependent protein kinases, PRX: peroxidases.

## MATERIALS AND METHODS

### Preparation and characterization of CDs

The CDs were synthesized by hydrothermal method using citric acid and ethanolamine at a molar ratio of 50:1, as described in our previous study (Li et al., 2021). The obtained CDs were redissolved in ultrapure water (200 mg/mL) and stored at room temperature. The morphology and chemical composition of CDs were studied using the FEI Tecnai G2 F20 transmission electron microscope (FETEM), Thermo Scientific Nicolet iS20 FTIR spectrophotometer, Thermo Scientific K-Alpha X-ray photoelectron spectroscope (XPS), and Bruker Avance III HD 500MHz nuclear magnetic resonance spectroscopy (NMR). UV-Vis absorption, excitation, and emission spectra of the CDs were recorded using an F-7000 Hitachi fluorescence spectrophotometer and a Perkin Elmer UV–Vis spectrophotometer (Lambda 750).

### Rice seedlings cultivation and treatments of CDs and stresses

Rice seedlings (Huahang 31) were cultivated in hydroponics using a half-strength Hoagland nutrient solution. Rice seeds were immersed in 75% ethanol for 2 min, rinsed five times with deionized water, and germinated in a specific sprouting tray containing ultrapure water in the dark at 30 °C. After 5 days, the rice seedlings were acclimated to light in an incubator for 24 h. Subsequently, homogeneous rice seedlings were transplanted into pots (12 rice seedlings per pot) containing half-strength Hoagland nutrient solutions. All rice seedlings were grown in a greenhouse at 30 °C for 16 h under the light conditions of 400 μmol m^−2^·s^−1^ and 28 °C for 8 h in darkness. The nutrient solutions were refreshed once a week.

After transplantation, CDs solutions (0, 100, 200, and 400 μg/mL) were sprayed on rice seedlings once a day with 5 mL/pot for two weeks to measure the effect of spraying CDs on rice growth. To study the enhancement of pre-spraying of CDs to rice stress resistance, rice seedlings were pre-sprayed with CDs for one week and then exposed to NaCl at 100 mM, CdCl_2_ at 1.63 mg/L (Cd at 1 mg/L), and 2,4-D at 5 mg/L in nutrient solutions, respectively. The rice seedlings were cultivated for another week without CDs spraying. Rice seedlings grown in normal conditions were used as the control treatment. Each treatment was conducted with three repetitions. Rice seedlings sprayed with CDs (100 μg/mL) prior to the addition of stress were used to study the regulatory mechanism of CDs.

### Root phenotype, anatomical, and microscopic structures of rice seedlings

To analyze the changes in root phenotype induced by CDs, rice roots from both control and CDs (100 μg·mL^−1^) treatments were imaged using an LA-S root scanning system. The total length, surface area, volume, and average diameter of the roots were then analyzed using WinRHIZO software (Hangzhou Wanshen Detection Technology Co., Ltd., China). The fresh weights of rice seedlings were also measured. At least three biological replicates of both control and CDs treatments were collected for analysis. For the anatomical structures, root samples from control and CDs treatments were collected and immersed in FAA fixative solution for 24 h. Then, the root samples underwent dehydration, transparency, wax filling, embedding, slicing, dewaxing, and sarranine-fast green staining, and were photographed using an optical microscope in succession. Besides, Hitachi S-4800 scanning electron microscope (SEM) and Hitachi-7800 TEM were used to observe the microscopic structures of rice roots following a standard protocol.

### Vitality, cytoplasm content, and mineral nutrient element contents in rice root

Rice root vitality was assessed using 2,3,5-triphenyltetrazolium chloride (TTC) following an established method.(Comas et al., 2000) To measure cytoplasm contents, the isolated rice roots were submerged in ultrapure water and sonicated for 30 min in an ice bath. Cytoplasm content was measured as the conductivity of the solution per gram of fresh root. Rice roots from both the control and CDs treatment were measured with five biological replicates.

Mineral nutrient element contents in rice roots were measured using inductively coupled plasma mass spectrometry (ICP-MS, Agilent 7700X, USA). Briefly, rice roots from the control and CDs treatments were collected, dried in an oven, and ground into powders using a tissue grinder. Then, 0.1 g samples were added to Teflon digestion tubes with 10 mL of concentrated nitric acid and heated at 115 °C for 30 min in a microwave digestion system. After dilution, the levels of P, K, Ca, Mg, and Mn were quantified using ICP-MS. A blank and a standard solution were utilized during the measurement to verify precision. The recoveries of the standard solution ranged from 96.7% to 108.4% and relative standard deviation values of each sample lower than 3% were accepted for the resolution. Five biological replicates of each treatment were measured, and the element concentration was calculated as milligrams per gram (mg/g) of tissue dry mass.

### Transcriptomic analysis of rice seedlings

Six biological replicates of rice seedlings from both the control and CDs treatments were collected, frozen in liquid nitrogen, and stored at −80 °C. The total RNA of these rice seedlings was extracted using TRIzol^®^ Reagent according to the manufacturer’s instructions. Then, the quality of RNA samples was determined using the 5300 Bioanaluser (Agilent), and the concentration was quantified with the ND-2000 (NanoDrop Technologies). RNA samples with an OD260/280 ratio of 1.8~2.2, OD260/230≥2.0, RIN≥6.5, 28S:18S≥1.0, and mass≥1 μg were used to construct the sequencing library following the Illumina® Stranded mRNA Prep, Ligation Kit from Illumina (San Diego, CA). The expression level of each transcript was calculated using the transcripts per million reads (TPM). DESeq2 software was used to analyze the differentially expressed genes (DEGs) between control and CDs treatments. DEGs with |log2FC|≥1 and FDR<0.05 were considered significantly differentially expressed. The functional annotation and enrichment analysis of DEGs were conducted using the EggNOG, Gene Ontology (GO), and Kyoto Encyclopedia of Genes and Genomes (KEGG) databases. RNA-Seq results were further confirmed using the qRT-PCR method by measuring the expression levels of 13 selected genes (Methods S1, Table S1, and Figure S1).

### Metabolomic analysis of rice seedlings

Six biological replicates of whole rice seedlings from both the control and CDs treatments were collected and ground into powders with liquid nitrogen for the non-targeted metabolomic analysis. The metabolite was extracted by mixing 50 mg of each sample with 400 μL extracting solution (methyl alcohol: water=4: 1, v: v) that contained 0.02 mg/mL L-2-chlorophenylalanine and being ground for 6 min at −10 °C and 30 min at 5 °C. The supernatant was used for the measurement of ultra-high performance liquid chromatography-tandem Fourier transform mass spectrometry (Thermo UHPLC-Q Exactive HF-X). The data were analyzed through the majorbio cloud platform (cloud.majorbio.com). Differentially expressed metabolites (DEMs) were identified using variable importance in projection (VIP) values (VIP ≥1 and *p* < 0.01).

### Statistical analysis

All data were presented as means ± standard deviation (SD) in this paper. Statistical significance was analyzed using a one-way analysis of variance (ANOVA) and compared using Duncan’s test at *p* < 0.05 levels by SPSS 24.

## Supplementary data

The following materials are available in the online version of this article.

**Supplementary Method S1**. Quantitative reverse transcription polymerase chain reaction (qRT-PCR).

**Figure S1.** Correlation between RNA-Seq and qRT-PCR for thirteen selected genes.

**Figure S2**. The optical characteristics of CDs.

**Figure S3**. Effects of CDs on rice seedling growth by foliage spray.

**Figure S4**. Photographs of rice seedlings pre-sprayed with and without CDs under control and heavy Cd, salt, and herbicide 2.4-D stresses.

**Figure S5**. TEM images show the distribution of CDs in rice leaf and root.

**Figure S6**. Transcriptional changes in rice seedlings sprayed with CDs.

**Figure S7**. Gene expression levels of DEGs enriched in adaptability-related KEGG pathways in CDs-treated rice seedlings.

**Figure S8**. Influence of CDs on the antioxidant defense system of rice.

**Figure S9**. Metabolome changes in rice seedlings sprayed with CDs.

**Table S1.** List of primers used for qRT-PCR in this study.

**Table S2.** CDs-induced DEGs annotated in EggNOG functions of cell cycle control, cell division, chromosome partitioning, and cytoskeleton.

## Acknowledgments

This work was supported by the National Natural Science Foundation of China (Grant No. 42207032, 52070064); Advanced Talents Incubation Program of the Hebei University (521100222012); Science Research Project of Hebei Education Department (BJK2024161). We thank for the financial support from Collaborative Innovation Center for Baiyangdian Basin Ecological Protection and Beijing-Tianjin-Hebei Sustainable Development and Institute of Life Sciences and Green Development of Hebei University.

## Author contributions

Yadong Li conceived this project and wrote the manuscript. Ronghua Xu, Jingyi Qi, Shang Lei, Qianying Han, Yunlong Ru performed the experimental work, and collected, analyzed, and interpreted the data. Congli Ma interpreted and discussed the data. Hongjie Wang critically revised the manuscript and finally approved the version to be published.

## Data availability

The data supporting the findings of this study are available in the article and its supplementary information files.

